# Emergent modularity in large language models: Insights from aphasia simulations

**DOI:** 10.1101/2025.02.22.639416

**Authors:** Chengcheng Wang, Zhiyu Fan, Zaizhu Han, Yanchao Bi, Jixing Li

**Author notes:** co-corresponding: Zaizhu Han, Yanchao Bi. first (Chengcheng Wang). co-first.

## Abstract

Recent large language models (LLMs) have demonstrated remarkable proficiency in complex linguistic tasks and have been shown to share certain computational principles with human language processing. However, whether LLMs’ internal components perform distinct functions, like semantic and syntactic processing in human language systems, remains unclear. Here, we systematically disrupted components of LLMs to simulate the behavioral profiles of aphasia—a disorder characterized by specific language deficits resulting from brain injury. Our findings showed that lesioning specific components of LLMs could replicate behaviors characteristic of different aphasia subtypes. Notably, while semantic deficits as those observed in Wernicke’s and Conduction aphasia, were relatively straightforward to simulate, reproducing syntactic and lexical impairments, as seen in Broca’s and Anomic aphasia, proved more challenging. Together, these results highlight both parallels and discrepancies between emergent modularity in LLMs and the human language system, providing new insights into how information is represented and processed in artificial and biological intelligence.

## Introduction

Recent large language models (LLMs), such as GPT-4 (OpenAI et al., 2023),LLaMA-3 (Grattafiori et al., 2024) and DeepSeek-V3 (DeepSeek-AI et al., 2024), have achieved remarkable performance across a wide range of natural language processing (NLP) tasks. These models are thought to share certain computational principles with the human brain during language processing, inspiring a number of “model-brain alignment” studies (Caucheteux et al., 2023; Caucheteux & King, 2022; Gao et al., 2024; Gao et al., 2024; Goldstein et al., 2022; Kumar et al., 2024; Schrimpf et al., 2021; Toneva & Wehbe, 2019; Yu et al., 2024). Despite their impressive capabilities, a significant challenge persists: LLMs operate as “black boxes,” making it difficult to mechanistically interpret how their predictive capabilities align with human language processing. In particular, it remains unclear whether there are distinct modules within LLMs that correspond to subcomponents of human language systems, such as lexical, semantic, syntactic, and discourse-level processing.

While some recent neuroimaging studies have suggested overlapping brain regions for syntactic and semantic processing (e.g., Blank et al., 2016; Blank & Fedorenko, 2020; Fedorenko et al., 2012, 2020; Shain et al., 2024), mounting neuropsychological evidence from aphasia research suggests a more modular architecture of language processing (Dronkers & Ivanova, 2023; Hickok & Poeppel, 2000; Pylkkänen, 2019), where damage to one system can selectively impair specific linguistic functions while sparing others. Aphasia is defined as an acquired impairment in language production, comprehension, or repetition due to brain injury. Various subtypes of aphasia have been documented, each associated with distinct neural substrates and linguistic deficits. Among the major subtypes are Broca’s aphasia (Broca, 1861), Wernicke’s aphasia (Wernicke, 1874), Conduction aphasia (Lichteim, 1885), Anomic aphasia and Global aphasia. Broca’s aphasia is characterized by severe syntactic deficits, particularly in producing and comprehending complex sentence structures. It is typically associated with damage to the left posterior inferior frontal gyrus (LIFG) and frequently extends to the ventral precentral gyrus, lateral striatum, and surrounding white matter (Fridriksson et al., 2007, 2015). Wernicke’s aphasia results in fluent but meaningless speech, reflecting impaired semantic processing. The primary lesion site is the left posterior superior temporal gyrus (LpSTG), often extending into the posterior middle temporal gyrus and inferior parietal lobule (Dronkers et al., 2004; But see Matchin et al. (2022), Mesulam et al. (2015, 2019) for recent debates on the neural correlates of Wernicke’s aphasia). Conduction aphasia involves deficits in mapping form and meaning, characterized by frequent sound structure errors and self-correction attempts. It is traditionally linked to damage in the arcuate fasciculus, disrupting the connection between Wernicke’s and Broca’s areas (Palumbo et al., 1992). Anomic aphasia primarily affects lexical-semantic retrieval, where patients exhibit relatively preserved comprehension and production but struggle with word-finding difficulties. The lesion sites are variable across patients but commonly involve the inferior temporal and inferior parietal regions (Raymer et al., 1997). Global aphasia results in widespread impairment across lexical, semantic, and syntactic levels, with extensive damage to the entire left perisylvian cortex (Kemmerer, 2022, Ch2).

Despite substantial neuropsychological evidence supporting the modularity of the human language system, pretrained language models are typically treated as monolithic models (Qiu et al., 2024). One line of research has attempted to isolate the syntactic abilities of language models by designing targeted linguistic constructions and correlating model outputs with human behavioral data across different architectures (Asami & Sugawara, 2024; Linzen et al., 2016; Mueller & Linzen, 2023; Ryu & Lewis, 2021; Simoulin & Crabbé, 2021; Timkey & Linzen, 2023). However, such approaches often fail to disentangle other language modules, such as lexicon and semantics. Another method for investigating the functional specialization of language models involves simulating language disorders by lesioning specific model components (e.g., Dell et al., 1997; Farah & McClelland, 1991; Hinton & Shallice, 1991; McClelland & Elman, 1986; McClelland & Rogers, 2003; Plaut & Shallice, 1993; Rogers et al., 2004). However, these connectionist models were constrained by small-scale architectures and single-modality processing, ultimately failing to capture the complexity and interconnectedness of human language processing.

In this study, we utilize the multimodal LLM Visual-Chinese-LLaMA-Alpaca (VisualCLA; Cui et al., 2024; Yang et al., 2023) to perform a picture description task, a widely used diagnostic tool for assessing aphasia (Goodglass & Kaplan, 1983). VisualCLA integrates a vision encoder, a resampler for multimodal integration, and a fine-tuned LLaMA model (Touvron et al., 2023), enabling it to process both visual and textual inputs. As such, VisualCLA surpasses previous small-scale connectionist models, which lack the ability to perform multimodal tasks in an end-to-end manner akin to human processing. We systematically lesioned individual layers, self-attention heads, and critical parameters within the text model of VisualCLA, simulating language deficits analogous to those observed in human aphasia (see Fig. 1 for an overview of our analysis pipeline). By analyzing the post-lesion performance of the model and comparing it to behavioral data from individuals with different aphasia subtypes, we aim to identify functionally distinct modules within the LLM that parallel those in the human language system.

**Fig. 1.**
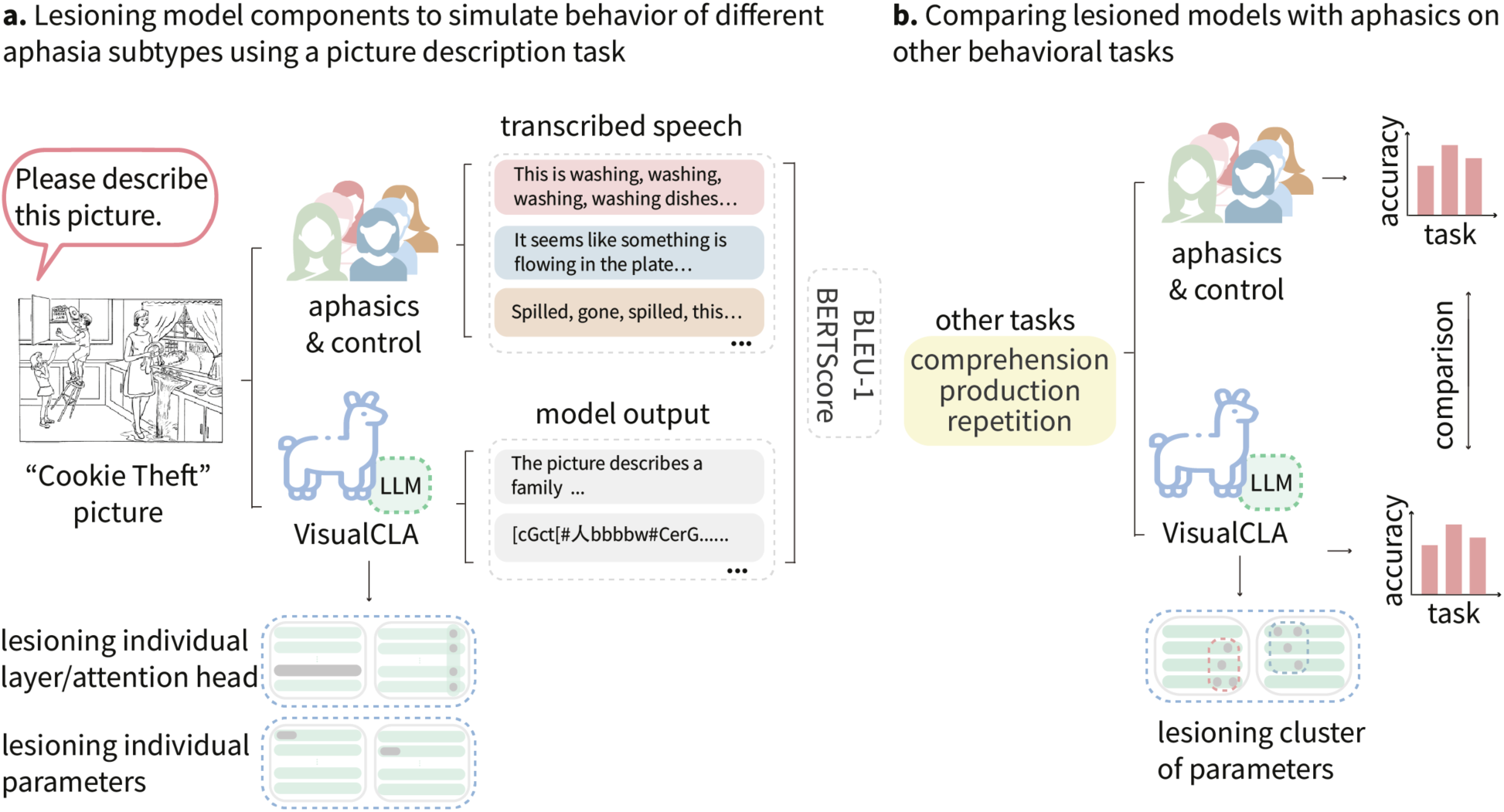
Overview of the analysis pipeline. **a**, Transcribed speech from the “Cookie Theft” picture description task was collected from five aphasia subtypes and control participants and compared to outputs from the VisualCLA model. The model was lesioned at the individual layer, self-attention head, and parameter levels. Model outputs were evaluated using BLEU-1 and BERTScore to quantify their similarity to aphasic speech. **b,** Clusters of lesioned parameters associated with each aphasia subtype were further tested on three additional behavioral tasks to assess comprehension, production, and repetition abilities. Accuracy scores from the lesioned models were compared with those of human participants, demonstrating the alignment between model-generated deficits and aphasic syndromes.

## Results

### Lesion map for each aphasia subtype and their corresponding functions

We used an existing dataset comprising 69 individuals with aphasia (17 females; mean age=46.9±12.1 years) from the China Rehabilitation Research Center and 43 healthy controls (21 females; mean age=49.3±10.7 years) from Beijing Normal University (Bi et al., 2015; Han et al., 2013). The patients were further categorized into 16 Broca’s aphasics, 11 Wernicke’s aphasics, 6 Conduction aphasics, 12 Anomic aphasics, and 24 Global aphasics. We first examined the overlaps of the lesion sites for different aphasia subtypes. We found that Broca’s aphasia showed the highest lesion overlap in the left frontal lobe; Wernicke’s aphasia showed the highest degree of overlap in the left temporal regions. Conduction aphasia was localized to the arcuate fasciculus and surrounding cortical areas, and Anomic aphasia was more diffusely distributed across the brain. Global aphasia lesions spanned extensive areas of the left hemisphere, covering both frontal and temporal regions, consistent with the severe language deficits observed in this condition (see Fig. 2a). These lesion sites are highly consistent with the typical neural correlates associated with different aphasia subtypes, as reported in the literature (e.g., Dronkers & Baldo, 2009; Kemmerer, 2022, Ch. 2; see Fig. 2b). According to the classical Wernicke–Lichtheim–Geschwind “house” model, Broca’s area and Wernicke’s area are primarily associated with language production and comprehension, respectively (see Fig. 2b, from Gazzaniga et al., 2009, p. 426). Additionally, these two regions play crucial roles in syntactic and semantic processing, with Broca’s area implicated in syntax and Wernicke’s area in semantics (Li et al., 2024; Li & Pylkkänen, 2021; Matchin & Hickok, 2020; Pylkkänen, 2019). fMRI term-based meta-analysis from Neurosynth (Yarkoni et al., 2011) further demonstrated a strong correspondence between Broca’s lesion sites and syntactic processing regions in the left inferior frontal gyrus (LIFG), as well as a high correspondence between Wernicke’s lesion sites and semantic processing regions in the left posterior superior temporal gyrus (LpSTG; see Fig. 2c).

**Fig. 2.**
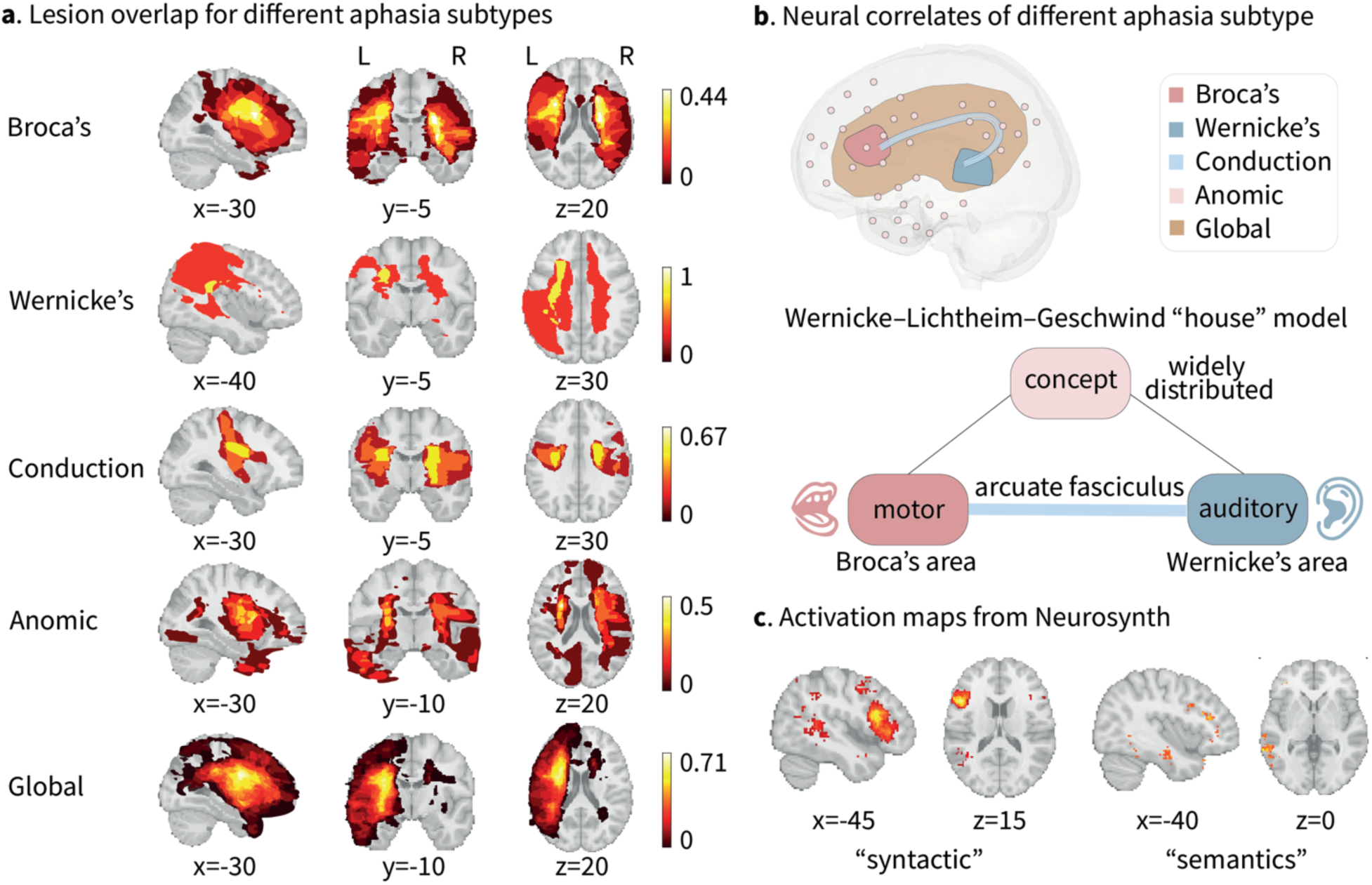
Lesion sites for different aphasia subtypes and their linguistic functions. **a,** Lesions overlap for each aphasia type overlaid on a standardized brain template. The lesioned voxels for each participant were assigned a value of 1, and the summed lesion overlaps were normalized by dividing by the number of patients within each aphasia group. **b,** Neural correlates of different aphasia subtypes and the classic Wernicke-Lichtheim-Geschwind “house” model for the neural architecture of language (from Gazzaniga et al., 2009, p. 426). **c,** Activation maps for the terms “syntactic” and “semantics” from Neurosynth, an fMRI term-based meta-analysis tool.

### Human and model performance on the picture description task

#### Syntactic features

We conducted an initial assessment of the behavioral outputs from the “Cookie Theft” picture description task, comparing responses from the intact VisualCLA model, the Control group, and different aphasia subtypes. We first calculated the average number of words and unique words per sentence, and we observed group-specific patterns consistent with previous literature (see Fig. 3a): Broca’s and Global aphasics produced shorter sentences with fewer unique words (Broca’s: 5.07±3.64 total and 4.5±3.05 unique words; Global: 4.21±3.45 total and 3.48±2.6 unique words), whereas other aphasia subtypes demonstrated a relatively preserved ability to produce longer and more complete sentences (Wernicke’s: 6.04±4.91 total and 5.55±4.2 unique words; Conduction: 7.56±5.17 total and 7±4.55 unique words; Anomic: 7.09±4.61 total and 6.46±4 unique words). The Control group and the intact VisualCLA model produced the longest sentences with most unique words (Control: 11.08±6.58 total and 10.04±5.68 unique words; VisualCLA: 10.25±4.51 total and 9.74±3.57 unique words). Supplementary Table 1 and Table 2 present the statistical results from a one-way analysis of variance (ANOVA) assessing group differences on the number of words and unique words per sentence, along with pairwise t-tests. No significant differences were observed between VisualCLA and the Control group, indicating that the model’s linguistic performance closely aligns with that of healthy individuals.

**Fig. 3.**
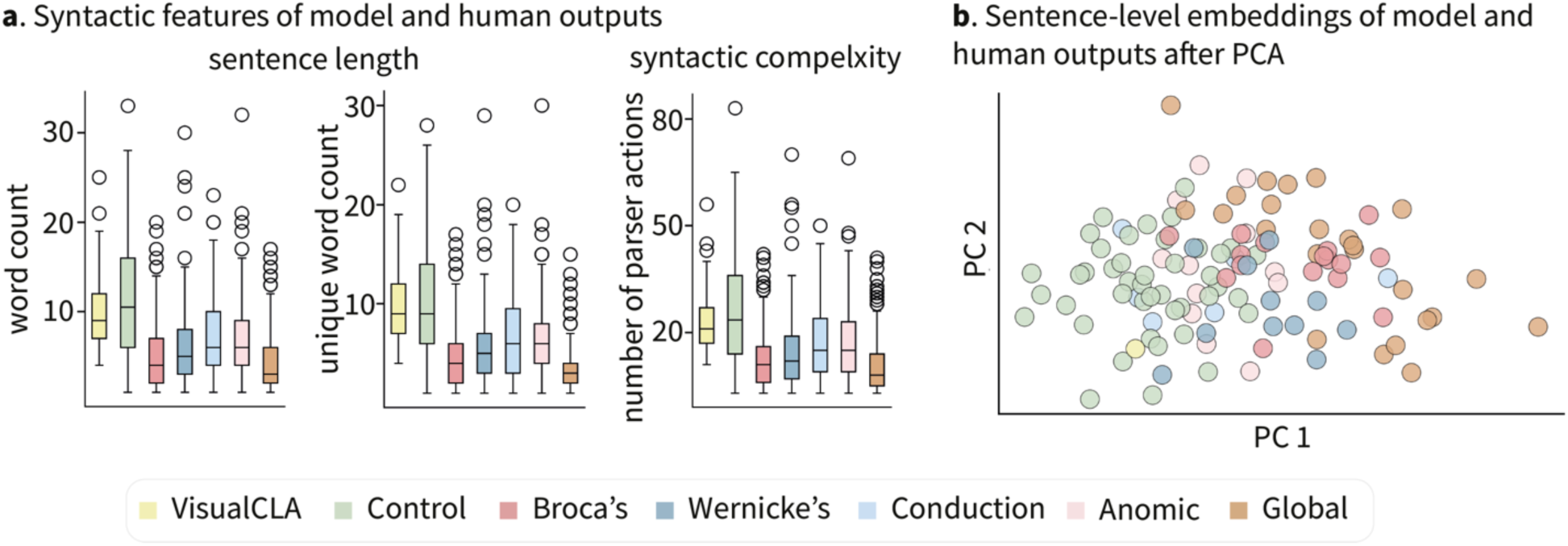
Syntactic and semantic characteristics of human and model outputs in the picture description task. **a,** Mean word count, unique word count and the total number of left-corner parsing steps per sentence across different aphasia subtypes, the control group, and the intact VisualCLA model. **b,** Sentence-level embeddings of human and model outputs, visualized after dimensionality reduction using PCA.

We also computed “syntactic complexity” of the aphasics and the Control group’s output using the total number of parser actions for each word within each sentence based on the left-corner parsing strategy. This complexity metric is associated with certain aspects of Yngve’s (1960) Depth hypothesis, where the processing effort required for a given word is based on its syntactic structure and a parsing strategy (Hale, 2014). Prior research has shown significant left temporal and frontal activity for the left-corner parsing strategies (Nelson et al., 2017), supporting it as a tentative model of how human subjects process sentence structures. Our results revealed group-specific patterns in left-corner parsing steps, with the LLM and the Control group demonstrating the highest mean number of parsing steps (VisualCLA: 22.97±8.63; Control: 25.42±14.53), indicating comparable syntactic complexity of their output. In contrast, outputs from Broca’s and Global aphasia exhibited significantly fewer parsing steps (Broca’s: 12.26±8.06; Global: 10.39±7.65), aligning with their hallmark fragmented speech and reduced syntactic complexity. Meanwhile, individuals with Wernicke’s, Conduction and Anomic aphasia (Wernicke’s: 14.68±11.05; Conduction: 18.17±11.85; Anomic: 16.82±10.37) showed an intermediate number of parsing steps, suggesting their relatively intact or partially preserved syntactic processing abilities (see Fig. 3a). Pairwise t-tests confirmed significant differences in the mean number of parser actions per sentence between Broca’s and Global aphasia and Wernicke’s, Conduction and Anomic aphasia (see Supplementary Table 3 for statistical results from ANOVA and pairwise t-tests).

#### Semantic features

For semantic features, we extracted sentence-level embeddings from the VisualCLA text model for both model-generated outputs and participant responses. Fig. 3b visualizes these embeddings after dimensionality reduction using Principal Component Analysis (PCA). The results suggest that VisualCLA closely parallels the Control group, whereas Broca’s and Global aphasia exhibit greater deviations, reflecting impaired descriptive language production. Additionally, Wernicke’s aphasia forms a distinct clustering pattern, separate from Broca’s aphasia, highlighting key differences in their linguistic deficits.

### Model performance after lesioning individual layers and self-attention heads

To further explore whether specific submodules of the model correspond to functional specialization within the human language network, we systematically lesioned the text model of VisualCLA at both the layer and self-attention head levels. Prior research has shown that the depth of a Transformer architecture (Vaswani, 2017) is critical for learning syntactic generalizations (Mueller & Linzen, 2023). Additionally, self-attention mechanisms have been shown to parallel cue-based retrieval theories of working memory in human sentence processing (Ryu & Lewis, 2021; Timkey & Linzen, 2023). We therefore hypothesize that lesioning layers within the model may lead to syntactic impairments resembling those observed in Broca’s aphasia. Lesioning attention heads may result in semantic deficits, similar to those characteristic of Wernicke’s aphasia.

The text model of VisualCLA is a fine-tuned Chinese LLaMA (Touvron et al., 2023) consisting of 32 layers, each with 32 attention heads. We separately lesioned each model layer and attention head, then generated responses to the “Cookie Theft” picture description task following the lesioning procedure. We employed BLEU-1 (Papineni et al., 2002) and BERTScore (T. Zhang et al., 2020) to evaluate the similarity between lesioned model outputs and aphasic speech. Our results showed that lesioning a single layer and self-attention head resulted in deficits more similar to Wernicke’s and Conduction aphasics (see Fig. 4 for the BLEU-1 and BERTScores comparing model outputs to outputs from all aphasia subtypes; statistical results from non-parametric t-tests are shown in Supplementary Table 4 and 5). Since neither Wernicke’s nor Conduction aphasia is marked by severe syntactic impairments, our initial hypothesis—that model layers play a more critical role in syntactic processing while self-attention primarily supports semantic processing— was not supported. It might be the case that syntactic functions in a Transformer architecture are more distributed rather than being localized within specific layers or attention heads. To further explore this possibility, we conducted additional analyses to evaluate the functional contributions of individual parameters within the model.

**Fig. 4.**
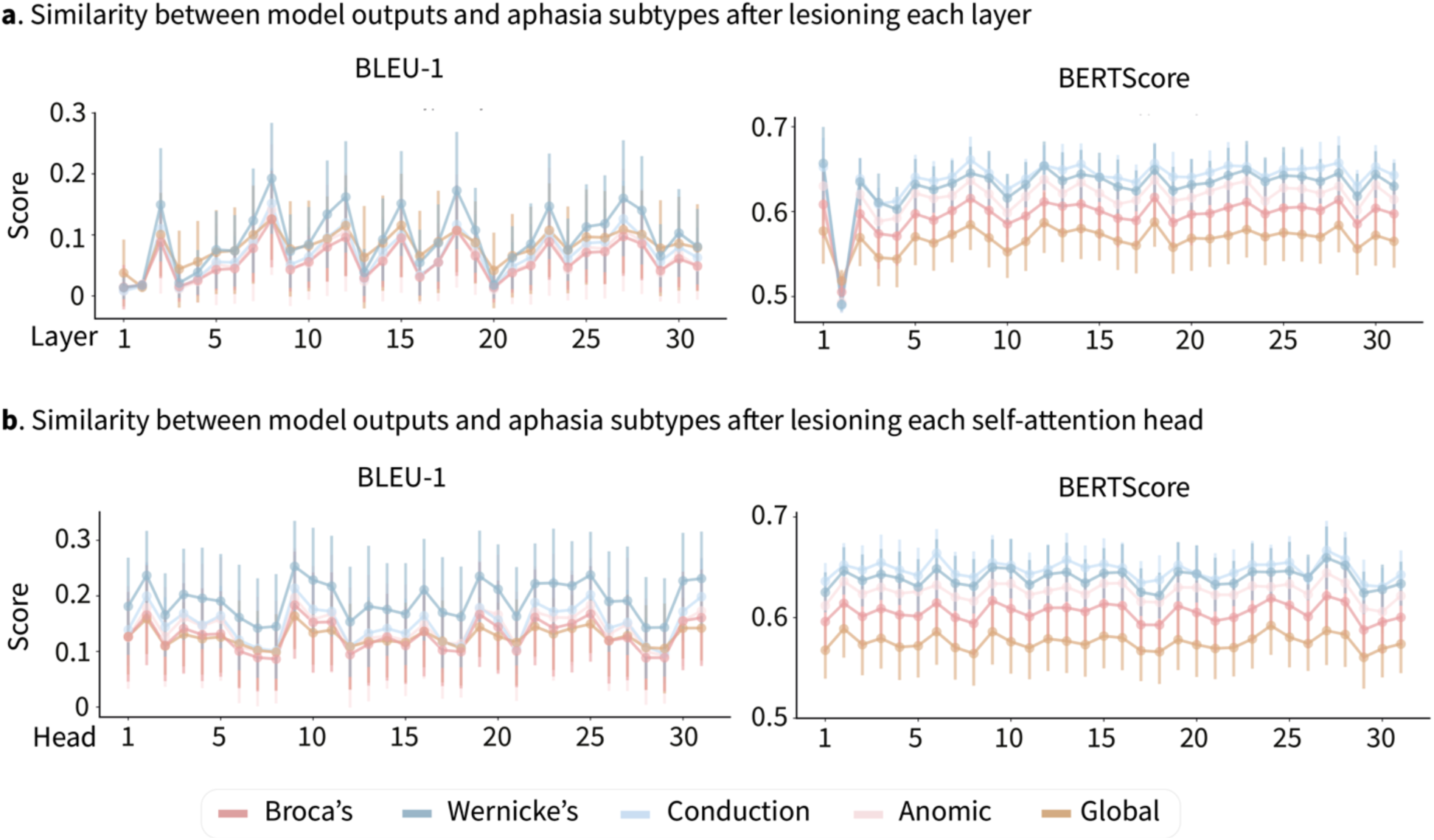
Similarity between model outputs and aphasic speech after lesioning individual layers and self-attention heads. **a,** BLEU-1 and BERTScore evaluated on outputs from each aphasia subtype after lesioning individual layers. **b,** BLEU-1 and BERTScore evaluated on outputs from each aphasia subtype after lesioning individual self-attention heads.

### Model performance after lesioning individual parameters

We fine-tuned the text model of VisualCLA using outputs from the Control group for the picture description task. We quantified the relative impact of each parameter by analyzing the magnitude of their gradient changes, following the methods outlined by Zhang et al. (2024). The text model of VisualCLA consists of 32 layers, each containing 7 submodules—4 attention blocks and 3 feedforward blocks—resulting in a total of 224 submodules. Within each submodule, we identified the top 1% of parameters exhibiting the greatest gradient changes. These high-impact parameters were subsequently lesioned, and the model’s outputs were collected for the “Cookie Theft” task. The resulting 224 outputs were compared to human responses across six different aphasia types using BLEU-1 and BERTScores. To determine each submodule’s associated aphasia subtype, we averaged the standardized BLEU-1 and BERTScores for each type and assigned the subtype with the highest scores to that submodule. Since lesioning only the top 1% of parameters from a single submodule was insufficient to fully reproduce any specific aphasic behavior, we iteratively grouped submodules into clusters and lesioned them collectively. For example, if two submodules showed higher BLEU-1 and BERTScore values for Broca’s aphasia, they were lesioned together, and the resulting outputs were reassessed. This process was repeated iteratively until lesioning a sufficient number of parameters successfully reproduced the targeted aphasic behavior.

Among the 224 submodules, lesioning the top 1% of high-impact parameters in 16 submodules produced deficits resembling Broca’s aphasia, while 5 submodules corresponded to Wernicke’s aphasia, 4 to Conduction aphasia, 15 to Anomic aphasia, and 3 to Global aphasia (see Fig. 5a). A larger number of submodules required for a given aphasia subtype suggests greater difficulty in reproducing that specific deficit, as more parameters in the model needed to be lesioned (see Supplementary Table 6 for the distribution of parameters identified as critical for each aphasia behavior across submodules). The lesioned models consistently exhibited symptoms characteristic of their respective aphasia clusters. For example, lesioning the cluster of parameters associated with Broca’s aphasia resulted in outputs characterized by simplified syntactic structures and frequent omissions. In contrast, lesioning parameters associated with Wernicke’s aphasia led to outputs exhibiting comprehension errors. Similarly, lesioning parameters associated with Global aphasia produced highly incoherent outputs, closely resembling the severe language impairments observed in human participants with this condition. Fig. 5b presents representative examples of model outputs after lesioning each parameter cluster, alongside each aphasia subtype’s performance on the “Cookie Theft” picture description task (see Supplementary Table 7 for five additional example outputs from each lesioned model).

**Fig. 5.**
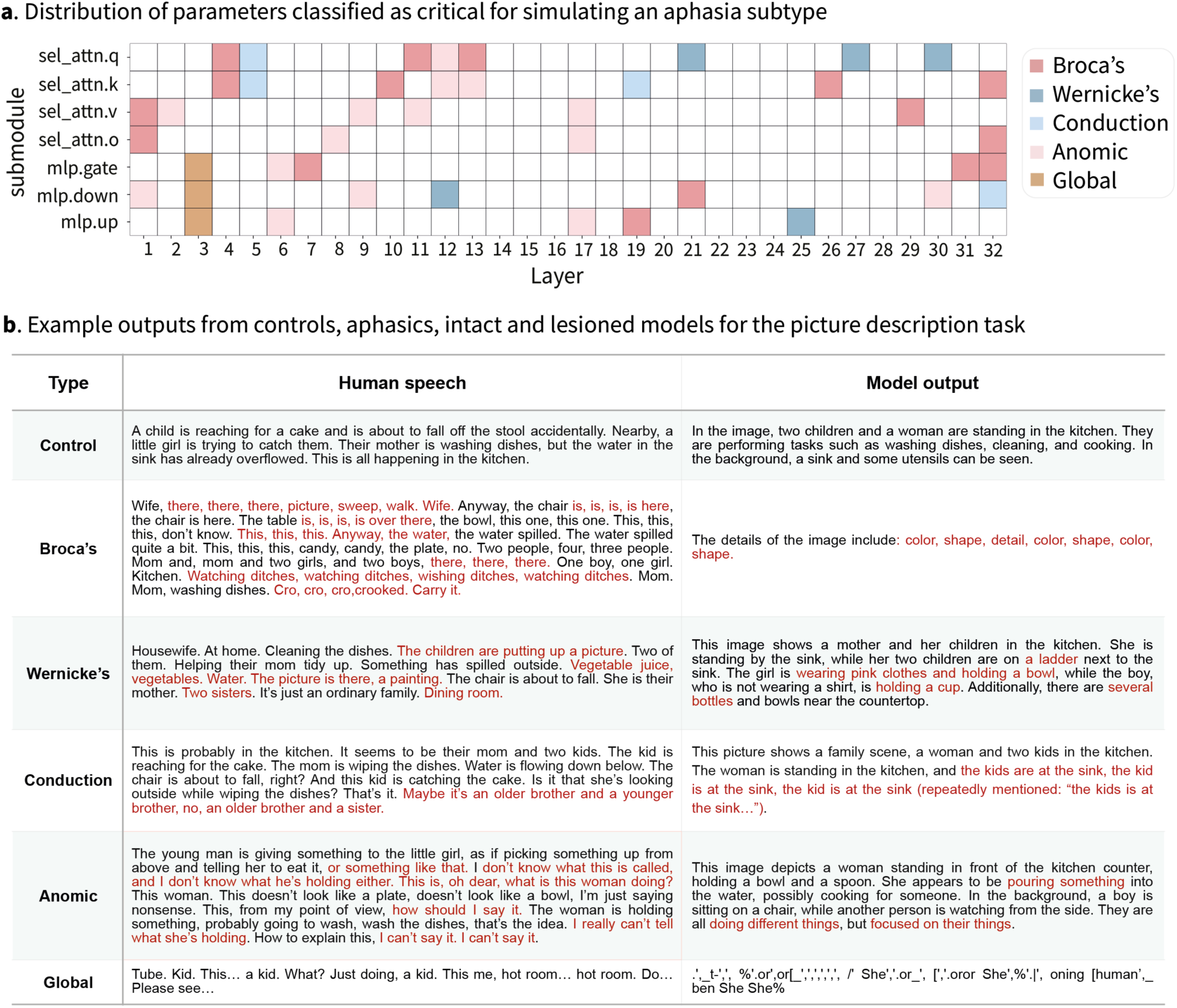
Results after lesioning high-impact parameters within each submodule of the VisualCLA text model. **a,** Distribution of parameters identified as critical for simulating behaviors associated with different aphasia subtypes. Each square represents the top 1% of parameters within a submodule that exhibited the greatest gradient change during fine-tuning. **b,** Example outputs from control participants, aphasic individuals, the intact VisualCLA model, and lesioned models on the “Cookie Theft” picture description task.

Contrary to the extensive lesion size typically associated with Global aphasia in human cases, we found that it was the easiest to simulate in lesioned models, requiring the lesioning of only the top 1% of parameters from three submodules. A closer examination of the model’s output suggests that these parameters may be crucial for encoding Chinese characters, as their removal resulted in the generation of random symbols such as ‘t-000}’. In contrast, Broca’s aphasia, the most common aphasia subtype in human cases, was the most challenging to replicate in lesioned models, requiring lesioning parameters from 16 submodules. Wernicke’s and Conduction aphasia were relatively easier to simulate, requiring lesioning top 1% of parameters from only 5 and 4 submodules, respectively. Anomic aphasia was also difficult to reproduce, as it involved lesioning parameters from 15 submodules.

### Performance of lesioned models on other behavioral tasks

To further validate that the lesioned models effectively parallel different aphasia subtypes, we evaluated their performance on three additional behavioral tasks: word associate matching, oral picture naming and oral word repetition. These tasks were designed to assess comprehension, production, and repetition abilities—key diagnostic criteria for classifying classic aphasia subtypes. Our results showed that accuracy scores of each lesioned model on the three behavioral tasks aligned more closely with their respective human aphasia subtypes (see Fig. 6a,b). Specifically, the model simulating Broca’s aphasia performed worse on oral picture naming and oral word repetition. The model simulating Wernicke’s aphasia struggled with word associate matching as well as oral word repetition. The Conduction aphasia model performed relatively well on comprehension and production tasks but exhibit deficits in oral word repetition. The Anomic aphasia model showed impairments in word associate matching but not in oral word repetition. Finally, the Global aphasia model performed the worst across all tasks. The findings further support the functional specificity of the identified parameters within the LLM (see Supplementary Table 8 on the accuracy scores of each aphasia subtype and lesioned model for each behavioral task).

**Fig. 6.**
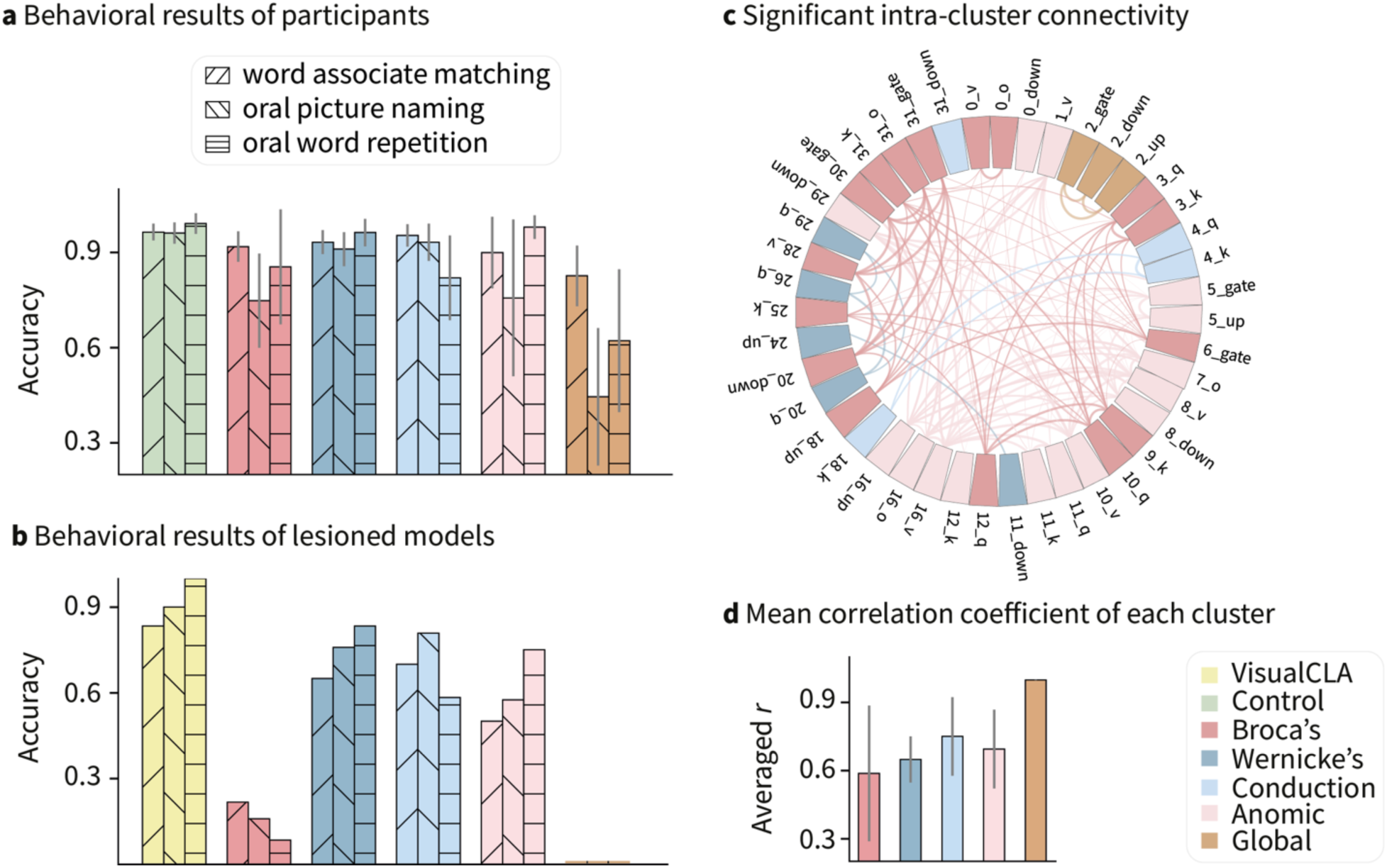
Results of lesioned models on other behavioral tasks and connectivity within clusters of parameters. **a,** Accuracy scores of aphasiacs and healthy controls on 3 additional behavioral tasks: word associate matching, picture associate matching, oral picture naming, and oral word repetition. **b,** Accuracy scores of each lesioned model on the 3 tasks. **c,** Statistically significant positive intra-cluster connectivity of top 1% parameters in the text model of VisualCLA. The color of the cells represents the cluster to which the parameters belong, while the intensity of the lines. **d,** Mean correlation coefficient of each cluster of parameters.

### Functional connectivity within clusters

The identified parameters associated with each aphasia subtype appear to be distributed throughout the model architecture (see Fig. 6c). This contrasts with human aphasia cases, where lesion sites are typically localized—for example, Broca’s aphasia is predominantly linked to lesions in the left inferior frontal gyrus (LIFG) rather than neuronal disruptions dispersed across the brain. To investigate the relationships among parameters classified under the same aphasia subtype, we examined the correlations of their gradient changes during fine-tuning the text model of VisualCLA with sentences from the Control group. This approach is analogous to functional connectivity analyses in brain research. Fig. 6c visualizes the connectivity networks for Broca’s, Wernicke’s, Conduction, Anomic, and Global aphasia clusters, where significant positive correlations are represented as edges linking the nodes (see Supplementary Table 9 for statistical results from non-parametric t-tests on the correlation coefficients between gradient changes of parameters). Global aphasia exhibited the strongest inter-cluster connectivity (r=0.998±0.001), indicating highly uniform parameter interactions within this group. Conduction aphasia showed the second-highest connectivity (r=0.75±0.17), followed by Anomic (r=0.7±0.17) and Wernicke’s aphasia (r= 0.65±0.1). Broca’s aphasia exhibited the lowest inter-cluster connectivity (r=0.58±0.29; see Fig. 6d). These findings further indicate that the parameters associated with syntactic processing, which are critical for inducing Broca’s aphasia, are less functionally connected, highlighting their distributed nature within the network.

## Discussion

In this study, we systematically lesioned components of an LLM and compared its behavioral deficits to aphasia subtypes, including Broca’s, Wernicke’s, Conduction, Anomic, and Global aphasia. We found that while semantic deficits as seen in Wernicke’s and Conduction aphasia were relatively easy to induce, syntactic and lexical impairments as seen in Broca’s and Anomic aphasia were harder to reproduce. This discrepancy likely arises from how information is represented and processed within LLMs, as well as the way they are trained to perform language tasks. Wernicke’s and Conduction aphasia are primarily characterized by comprehension deficits and disruptions in the mapping between comprehension and production. In LLMs, these deficits manifest as sequence generation errors, resulting in fluent but semantically incoherent outputs. One possible explanation is that LLMs maintain internal coherence by balancing text encoding (understanding context) and decoding (generating text). Lesioning critical parameters may disrupt the information flow between these processes, leading to fluent but contextually irrelevant responses, akin to the “word salad” phenomenon observed in Wernicke’s aphasia. Additionally, during text generation, LLMs select the most probable next word based on token probabilities. When critical weights in the prediction layers are lesioned, the model may lose its ability to constrain word choice based on contextual meaning, resulting in random yet syntactically plausible outputs. This pattern resembles the incorrect word substitutions and frequent self-corrections characteristic of Conduction aphasia.

In contrast, LLMs do not typically produce very short or syntactically incorrect sentences, as seen in Broca’s aphasia, nor do they exhibit word-finding difficulties characteristic of Anomic aphasia. This may be because LLMs are predominantly trained on massive corpora containing grammatically correct sentences from structured data sources, such as books, articles, and dialogues. This exposure leads to a strong bias toward syntactic correctness, making it difficult for lesions to induce syntactic breakdowns or lexical retrieval failures. Even when key semantic or lexical processing parameters are disrupted, the autoregressive nature of LLMs allows them to continue generating plausible syntactic structures. Moreover, our dataset consists solely of Chinese, a language that may exhibit greater flexibility in certain grammatical aspects, such as word order, compared to many Indo-European languages. Future investigations using diverse language samples would offer further insights into how syntactic disruptions manifest in lesioned LLMs across linguistic contexts.

The observation that LLMs exhibit emergent modularity presents an intriguing parallel to Fodor’s (1983) theory of modularity in the human mind. While LLMs were not explicitly designed with modularity in mind, our lesioning approach revealed that certain model parameters function in ways that resemble the human language system, particularly in comprehension processing and the mapping between comprehension and production. These findings suggest the potential for designing modular LLMs that map distinct linguistic functions to specific model components, enhancing both their interpretability and cognitive alignment. A modular architecture could foster greater parallels with human cognition, enabling more direct comparisons between artificial and biological language processing.

One potential limitation of this work is that computational lesioning may produce more severe deficits than those found in clinical populations, where redundancy and plasticity mitigate the impact of focal damage (Murphy & Corbett, 2009). For example, disabling a single “critical” subset of parameters within the model often yielded outputs resembling Global aphasia, a rare and extremely debilitating condition in humans (Hillis, 2007). However, biological brains can redistribute functions after injury (Dancause et al., 2005) in ways that artificial networks generally do not. Future work could approximate neural plasticity by implementing adaptive mechanisms— such as dynamic reweighting—to see whether the model can “recover” partial language functionality post-lesion.

To sum up, our study reveals that lesioning LLMs can simulate aphasia-like deficits, shedding light on the emergent modularity of artificial language systems. While semantic impairments were relatively easy to reproduce, syntactic and lexical deficits proved more challenging, highlighting key differences between biological and artificial language systems. The parallels between LLM lesioning and aphasia syndrome suggest that future models could benefit from more explicitly modular architectures, improving both their interpretability and cognitive plausibility.

## Methods

### Participants

We utilized an existing dataset comprising 69 individuals with aphasia (17 females; mean age=46.9±12.1 years; mean education level=12.8±3.7 years) from the China Rehabilitation Research Center and 43 healthy controls (21 females; mean age=49.3±10.7 years; mean education=13.7±3.8 years) from Beijing Normal University (Bi et al., 2015; Han et al., 2013). All participants were right-handed native Mandarin speakers. Among the 69 patients, 59 had experienced a stroke (cerebrovascular event), 9 had sustained traumatic brain injuries, and 1 had carbon monoxide poisoning. Based on the Aphasia Battery of Chinese (S. R. Gao et al., 1993), the patients were further categorized into 16 cases of Broca’s aphasia, 11 of Wernicke’s aphasia, 6 of Conduction aphasia, 12 of Anomic, and 24 of Global aphasia. Except for one individual with multiple strokes, each patient had a single injury, with the most recent event occurring approximately 3–4 months prior to the assessment (145.7±156.7 days). All participants were screened to ensure adequate post-stroke vision and hearing.

### Structural MRI data

The MRI data were collected using a 1.5 T General Electric SIGNATM Excite scanner at the China Rehabilitation Research Centre. Anatomical scans were obtained using a Magnetization Prepared RApid Gradient-Echo (MP-RAGE) sagittal sequence (248 single-shot interleaved sagittal slices; voxel size=0.49×0.49×0.70 mm; FOV=250 mm; TR=12.26 ms; TE=4.2 ms; TI=400 ms; flip angle=15°, slice number=248 slices). FLAIR T2-weighted images were acquired using an axial sequence (28 single-shot interleaved axial slices; voxel size =0.49×0.49×5 mm; FOV=250 mm; TR=8002 ms; TE=127.57 ms; TI=2000 ms; flip angle=9°). The T1-weighted MRI scans were acquired twice for quality assurance, whereas the T2-weighted FLAIR images were acquired once. All structural MRI data were co-registered using a trilinear interpolation method in SPM5 and averaged. T2-weighted FLAIR images were co-registered and resliced to the native space of the averaged 3D images. Structural images were resliced (voxel size=1×1×1 mm) and normalized to Montreal Neurological Institute (MNI) space for group-level analyses (see Bi et al., 2015 and Han et al., 2013 for detailed description of the MRI acquisition and preprocessing procedures). Lesion contours for each participants were manually delineated slice by slice by two trained researchers, with reference to T2 FLAIR images, and verified by an experienced radiologist. The lesioned voxels for each participant were assigned a value of 1, and the summed lesion overlaps were normalized by dividing by the number of patients within each aphasia group. We also extracted the activation map for the terms “syntactic” (169 studies) and “semantics” (84 studies) from fMRI term-based meta-analysis from Neurosynth (Yarkoni et al., 2011).

### Behavioral tasks

Apart from the “Cookie Theft” picture description task, we selected 3 additional tasks from the 32 behavioral tasks originally conducted by Bi et al. (2015) and Han et al. (2013): the word associate matching task, the oral word naming task, and the oral word repetition task. These tasks were chosen for their ability to provide a comprehensive assessment of language comprehension, production, and repetition in both humans and LLMs. The selected tasks included a total of 60 trials for word associate matching, 120 trials for picture naming, and 12 trials for word repetition. These words in these tasks spanned six semantic categories: actions (e.g., play piano), animals (e.g., elephant), common artifacts (e.g., umbrella), fruits and vegetables (e.g., fruit), large non-manipulable objects (e.g., well), and tools (e.g., axe). We excluded celebrity faces from the trials because VisualCLA cannot identify celebrities. Participant’s oral responses were recorded by a portable digital recorder (SONY^TM^) and accuracy for the all the tasks (except for picture description) was scored on a binary scale (1 for correct, 0 for incorrect) by trained native Mandarin speakers for each trial across the four tasks.

In the **picture description** task, participants viewed the black-and-white “Cookie Theft” picture (see Fig.1) from the Diagnostic Aphasia Examination (BDAE; Goodglass & Kaplan, 1983) and were asked to describe the contents of the picture. No time limit was imposed. In the **word associate matching** task, participants were presented with three written words arranged in an upright triangle on a touchscreen. They were required to determine which of the two bottom words (e.g., penguin and elephant) was semantically closer to the top word (e.g., polar bear) by touching the corresponding word displayed on the touchscreen. Participants had a maximum of 60 seconds to complete each trial. In the **oral picture naming** task, participants were presented with colored images of objects (e.g., a yellow potato) on a computer screen and were asked to name each object aloud. The **oral word repetition** task includes eight words (e.g., “wrong”) and four sentences (e.g., “The teacher helps the children with their homework”). Participants listened to each stimulus and were instructed to repeat it aloud immediately after hearing it. Participants’ oral responses were manually transcribed and verified by multiple native Chinese speakers. Each word in the transcribed speech was annotated for part-of-speech (POS) using spaCy’s Chinese pipeline (Honnibal et al., 2020).

### Extracting syntactic features

From the speech transcripts generated by participants for the “Cookie Theft” picture description task, we calculated the total number of words and unique words per sentence for each participant, as well as for 10 outputs generated by the intact VisualCLA model (Cui et al., 2024; Yang et al., 2023). We performed a one-way analysis of variance (ANOVA) and pairwise two-sample t-tests to compare the mean number of words and unique words per sentence across the six participant groups (Broca’s, Wernicke’s, Conduction, Anomic, Global, Control) and the VisualCLA- generated outputs.

We also calculated the total number of parser actions per sentence from the output of each participant and the intact VisualCLA model based on context-free grammar (CFG) syntactic trees generated by the Stanford Parser (Levy & Manning, 2003). We applied the left-corner parser strategy which integrates elements of both top-down and bottom-up approaches, applying a grammatical rule upon encountering the first symbol on the right-hand side of the rule (Hale, 2014). The same analyses were conducted on 20 outputs generated by the VisualCLA model. To evaluate statistical significance, we performed an ANOVA and pairwise two-sample t-tests to compare number of parser actions across the six participant groups and the VisualCLA-generated outputs.

### Extracting semantic features

We computed sentence-level embeddings for outputs from each participant and the intact VisualCLA model by averaging token embeddings for each sentence from the text model of VisualCLA. Since the 10 outputs of the model were highly similar in meaning, we selected only one representative output for analysis. The text model of VisualCLA consists of 32 layers, and we selected embeddings from the 20th layer, as prior research suggests that activations at approximately two-thirds of the total layers most closely align with brain activity during language processing (Caucheteux & King, 2022). To investigate group differences in sentence meaning, we applied Principal Component Analysis (PCA) to the extracted embeddings, enabling a comparative analysis of semantic representations across groups.

### Simulating aphasic behavior by lesioning model components

To simulate aphasic behaviours, we systematically disabled specific components of the text model in VisualCLA, including individual layers, individual attention heads, or parameters from specific submodules. We then provided the lesioned models with the “Cookie Theft” image along with the text prompt “Please describe this picture.” in Chinese, setting the maximum token limit to 200 to encourage longer outputs. We analyzed whether these lesioned models exhibited language deficits analogous to recognized aphasia subtypes. This approach extends classic lesioning studies on connectionist models (e.g., Farah & McClelland, 1991; Plaut & Shallice, 1993) by applying them at scale in a multimodal LLM capable of performing the same picture-description task as humans.

### Lesioning individual layers and self-attention heads

The text model of VisualCLA consists of 32 layers (excluding the embedding layer), each containing 32 self-attention heads. We systematically disabled individual layers or attention heads and analyzed their impact on model performance during the “Cookie Theft” picture description task. For layer lesioning, we deactivated one entire layer at a time by setting all its parameters, including attention weights and feedforward sub-layers, to zero. This procedure resulted in 32 distinct lesioned models, each with a specific layer removed. For attention head lesioning, we disabled a single attention head at a time at the same positional index across all 32 layers. This approach produced another set of 32 lesioned models, each with one attention head removed.

For each model output after lesioning, we assessed its similarity to aphasic outputs using BLEU-1 (Papineni et al., 2002) and BERTScore (T. Zhang et al., 2020). BLEU-1 prioritizes exact word matches, measuring precision by counting unigram overlaps between the predicted and reference outputs. BERTScore evaluates semantic similarity, comparing word embeddings in the predicted and reference sequences.

### Lesioning individual parameters

In addition to lesioning individual model layers and self-attention heads, we also lesioned individual parameters within each submodule of the text model of VisualCLA. Specifically, we fine-tuned the model using outputs from the Control group and assessed the relative impact of each parameter by analyzing the magnitude of their gradient changes, following the methodology outlined by Zhang et al. (2024). Instead of exhaustively zeroing out every parameter and re-evaluating the model—a computationally prohibitive process—we used a first-order Taylor approximation to estimate parameter importance. This approximation calculated the absolute value of the product of a parameter’s value and its gradient during pre-training. By focusing on absolute values, we emphasized the magnitude of each parameter’s contribution to language processing, regardless of its direction of influence.

Formally, given a large corpus 𝒟 and model parameters 𝜃 = [𝜃_1_, 𝜃_2_, …, 𝜃*_d_*] ∈ *R^d^* where each 𝜃*_j_* ∈ 𝑅 represents the 𝑗-th parameter, the training objective is to minimize the loss function ℒ(𝒟, 𝜃) : ℒ(𝒟, 𝜃) = ∑*_x_*_∈𝒟_ ∑*_i_* log 𝑝_𝜃_ (𝑥*_i_*|𝑥_1_, …, 𝑥_i-1_), where 𝑥 = {𝑥_1_, …, 𝑥*_n_*} denotes an input token sequence and 𝜃 denotes parameters of the model. The importance of each parameter is denoted by ℐ(𝜃) ∈ 𝑅*^d^*, with ℐ_𝒿_(𝜃) representing its significance. Assuming an independent and identically distributed (i. i. d.) data setting, the importance of a parameter ℐ_𝒿_(θ) is quantified by the increase in prediction loss upon removing θ_j_. This is computed as the absolute difference between prediction losses with and without 𝜃*_j_* : ℐ_𝒿_(θ) = |ℒ(𝒟, θ) − ℒ(𝒟, θ|θ*_j_* = 0)|. Since directly computing ℐ_𝒿_(θ)for each parameter is computationally expensive, requiring 𝑑 separate evaluations of the model, each omitting a single parameter. This complexity escalates as the number of parameters, 𝑑, reaches hundreds of billions. To address this, we use a first-order Taylor expansion of 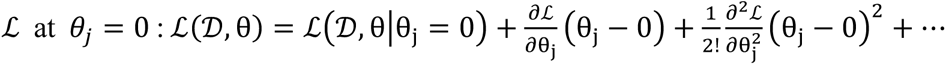 By approximating I_j_(θ) using only the first-order term, we eliminate the need for computing 𝑑 separate models: ℐ_𝒿_(θ) ≈ |g_j_θ_j_|, where 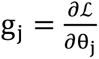 is the gradient of the loss with respect to θ_j_. Since gradients are readily available through backpropagation, this provides an efficient means of estimating parameter importance (see Zhang et al., 2024).

The text model of VisualCLA is a Chinese version of Alpaca-7B (Taori et al., 2023) consists of 32 layers, each containing 7 submodules (4 attention blocks and 3 feedforward blocks), resulting in a total of 224 submodules. We focused on parameters from the attention and feedforward layers, as these are directly involved in transforming and attending to information within the model. Parameters from the embedding layer (“embed_tokens”), normalization layers (“input_layernorm” and “post_attention_layernorm”), and the language model head (“lm_head”) were excluded from the analysis. To mitigate the potential confounding effects of parameters with massive activations, we implemented an additional filtering step following the methodology of Sun et al. (2024). Parameters with activation magnitudes exceeding predefined thresholds— indicative of massive activations functioning as fixed biases rather than dynamic components of language processing—were excluded. This ensured the analysis concentrated on meaningful variations relevant to the model’s linguistic capabilities. After filtering, we identified the top 1% of parameters for each of the 224 submodules. Each of these top-performing parameters was lesioned, and the model’s outputs were collected for the “Cookie Theft” picture description task. These 224 outputs were then compared against human responses across six different aphasic types to assess their alignment with specific language impairments.

To identify the aphasia subtype associated with each submodule, we calculated the average of the BLEU-1 and BERTScore metrics for each subtype and assigned the subtype with the highest average score to the corresponding submodule. Since lesioning only the top 1% of parameters within a single submodule did not fully replicate any specific aphasic behavior, we adopted an iterative approach by grouping submodules into clusters and lesioning them collectively. For example, if two submodules showed higher BLEU-1 and BERTScore values for Broca’s aphasia, they were lesioned together, and the resulting outputs were reassessed. This process was repeated iteratively until lesioning a sufficient number of parameters successfully reproduced the targeted aphasic behavior. All computational experiments are performed on a high-performance computing (HPC) cluster with 112 AMD EPYC 7522 CPUs and 512 GB ROM, and 8 NVIDIA A100-SXM4-80GB. Calculating the impact of each parameter in the text model of VisualCLA requires around 2 GPU hours.

### Validating lesioned models on other behavioral tasks

To validate the alignment of the lesioned models with their designated aphasic subtypes, we evaluated them on three other behavioral tasks: word associate matching, oral picture naming, and oral word repetition. Binary scores (1 for correct, 0 for incorrect) were assigned by two independent raters for each task. Inter-rater reliability calculated using Cohen’s Kappa (Cohen, 1960) reached 0.98, indicating high scoring consistency. The average score between the two human raters was used to represent the model’s accuracy for each trial. We compared the accuracy scores of the lesioned models on each behavioral task with the averaged accuracy scores of human participants across each aphasia subtype.

### Functional connectivity within clusters of parameters

To assess the relationship between each pair of the top 1% parameters within each attention and feedforward layer of the text model of VisualCLA, we further calculated pairwise correlations of the parameters’ weight changes during finetuning the model with the Control group’s output on the Cookie Theft description task. There are 336 sentences in total generated by the Control group, and for each sentence, we assessed the importance of individual parameters by calculating the prediction loss after removing each parameter, following Zhang et al (2024). We then averaged the gradient values across the 224 top 1% parameters for all 336 sentences, resulting in a 336*224 matrix. Each row corresponds to a sentence, and each column corresponds to a top 1% parameter. We selected the top 1% parameters that were identified for each aphasia subtype and calculated the pairwise Pearson’s correlation between these parameters across their 336 sentence-wise gradient values. This produced a 224*224 matrix of correlation coefficients. The statistical significance of the correlation coefficients was assessed by comparing the original r values for each pair of the top 1% parameters within each submodule to a null distribution generated by randomly shuffling the gradients and recalculating the correlations 10,000 times.

## Data availability

The aphasia dataset is available upon request.

## Code availability

All codes are available at https://github.com/compneurolinglab/aphasia

## Acknowledgements

This work was supported by the CityU Start-up Grant 7020086 and CityU Strategic Research Grant 7200747 (JL).

## Author contributions

JL designed the study; ZH and YB acquired the patient data; ZH analyzed the patient data; CW and ZF organized the patient data; CW and JL conducted modeling analyses; JL wrote the paper.

## Supplementary information

**Supplementary table 1.** Statistical comparisons of word count per sentence for the “Cookie Theft” task.

| **Test** | **Word count per sentence** |  |  |  |
| --- | --- | --- | --- | --- |
| ANOVA | ***F*** | ***p*** | **df1** | **df2** |
|  | 76.57 | 0.0000 | 6 | 1460 |
| *t*-test | **subtype 1** | **subtype 2** | ***t*** | ***p*** |
|  | Broca's | Wernicke's | -2.1152 | 0.0355 |
|  | Broca's | Conduction | -3.8444 | 0.0002 |
|  | Broca's | Anomic | -4.9339 | 0.0000 |
|  | Broca's | Global | 3.1436 | 0.0017 |
|  | Broca's | Control | -14.4701 | 0.0000 |
|  | Broca's | VisualCLA | -9.3891 | 0.0000 |
|  | Wernicke's | Conduction | -2.0667 | 0.0407 |
|  | Wernicke's | Global | 4.1201 | 0.0001 |
|  | Wernicke's | Control | -9.2823 | 0.0000 |
|  | Wernicke's | VisualCLA | -6.2856 | 0.0000 |
|  | Conduction | Anomic | 0.6747 | 0.5012 |
|  | Conduction | Global | 5.2535 | 0.0000 |
|  | Conduction | Control | -4.9532 | 0.0000 |
|  | Conduction | VisualCLA | -3.1396 | 0.0021 |
|  | Anomic | Global | 7.3109 | 0.0000 |
|  | Anomic | Control | -7.9637 | 0.0000 |
|  | Anomic | VisualCLA | -4.9045 | 0.0000 |
|  | Global | Control | -17.1323 | 0.0000 |
|  | Global | VisualCLA | -11.2698 | 0.0000 |

**Supplementary table 2.** Statistical comparisons of unique word count per sentence for the “Cookie Theft” task.

| **Test** | **Unique word count per sentence** |  |  |  |
| --- | --- | --- | --- | --- |
| ANOVA | ***F*** | ***p*** | **df1** | **df2** |
|  | 97.9707 | 0.0000 | 6 | 1460 |
| *t*-test | **subtype 1** | **subtype 2** | ***t*** | ***p*** |
|  | Broca's | Wernicke's | -2.7007 | 0.0075 |
|  | Broca's | Conduction | -4.4113 | 0.0000 |
|  | Broca's | Anomic | -5.5916 | 0.0000 |
|  | Broca's | Global | 4.5937 | 0.0000 |
|  | Broca's | Control | -15.5772 | 0.0000 |
|  | Broca's | VisualCLA | -11.3034 | 0.0000 |
|  | Wernicke's | Conduction | -2.2528 | 0.0259 |
|  | Wernicke's | Global | 5.5325 | 0.0000 |
|  | Wernicke's | Control | -9.6173 | 0.0000 |
|  | Wernicke's | VisualCLA | -7.5655 | 0.0000 |
|  | Conduction | Global | 6.3259 | 0.0000 |
|  | Conduction | Control | -4.8870 | 0.0000 |
|  | Conduction | VisualCLA | -3.9711 | 0.0001 |
|  | Anomic | Global | 8.9548 | 0.0000 |
|  | Anomic | Control | -8.2635 | 0.0000 |
|  | Anomic | VisualCLA | -6.2413 | 0.0000 |
|  | Global | Control | -19.4035 | 0.0000 |
|  | Global | VisualCLA | -13.9010 | 0.0000 |

**Supplementary table 3.** Statistical comparisons of left-corner parser steps per sentence for the “Cookie Theft” task.

| Test | Number of left-corner parser actions per sentence |  |  |  |
| --- | --- | --- | --- | --- |
| ANOVA | *F* | *p* | df1 | df2 |
|  | 73.9631 | 0.0000 | 6 | 1460 |
| *t*-test | **subtype 1** | **subtype 2** | *t* | *p* |
|  | Broca's | Wernicke's | -2.3581 | 0.0192 |
|  | Broca's | Conduction | -3.9905 | 0.0001 |
|  | Broca's | Anomic | -4.9916 | 0.0000 |
|  | Broca's | Global | 3.0638 | 0.0023 |
|  | Broca's | Control | -14.3389 | 0.0000 |
|  | Broca's | VisualCLA | -9.4151 | 0.0000 |
|  | Wernicke's | Conduction | -2.0756 | 0.0399 |
|  | Wernicke's | Global | 4.2910 | 0.0000 |
|  | Wernicke's | Control | -8.8539 | 0.0000 |
|  | Wernicke's | VisualCLA | -5.9775 | 0.0000 |
|  | Conduction | Global | 5.3203 | 0.0000 |
|  | Conduction | Control | -4.4901 | 0.0000 |
|  | Conduction | VisualCLA | -2.7453 | 0.0069 |
|  | Anomic | Global | 7.2820 | 0.0000 |
|  | Anomic | Control | -7.6918 | 0.0000 |
|  | Anomic | VisualCLA | -4.7146 | 0.0000 |
|  | Global | Control | -16.9390 | 0.0000 |
|  | Global | VisualCLA | -11.3014 | 0.0000 |

**Supplementary table 4.** Results of non-parametric t-tests assessing the model’s similarity to various aphasia subtypes on the “Cookie Theft” task following lesions to individual layers.

| Lesioning individual layers |  |  |  |  |  |
| --- | --- | --- | --- | --- | --- |
| Metric | subtype 1 | subtype 2 | Layer | *t* | *p* |
| BLEU-1 | Wernicke's | Broca's | 12-13 | 1.9964 | 0.0688 |
|  |  |  | 19-20 | 1.8793 | 0.0762 |
|  |  |  | 26-29 | 1.8305 | 0.0543 |
| BERTScore | Wernicke's | Broca's | 3-32 | 2.7550 | 0.0048 |
|  |  | Global | 3-32 | 5.4017 | 0.0000 |
|  | Conduction | Broca's | 3-32 | 2.8580 | 0.0036 |
|  |  | Anomic | 28-30 | 1.9317 | 0.0548 |
|  |  | Global | 3-32 | 4.9548 | 0.0001 |
|  | Broca's | Global | 3-32 | 2.6875 | 0.0040 |

**Supplementary table 5.**
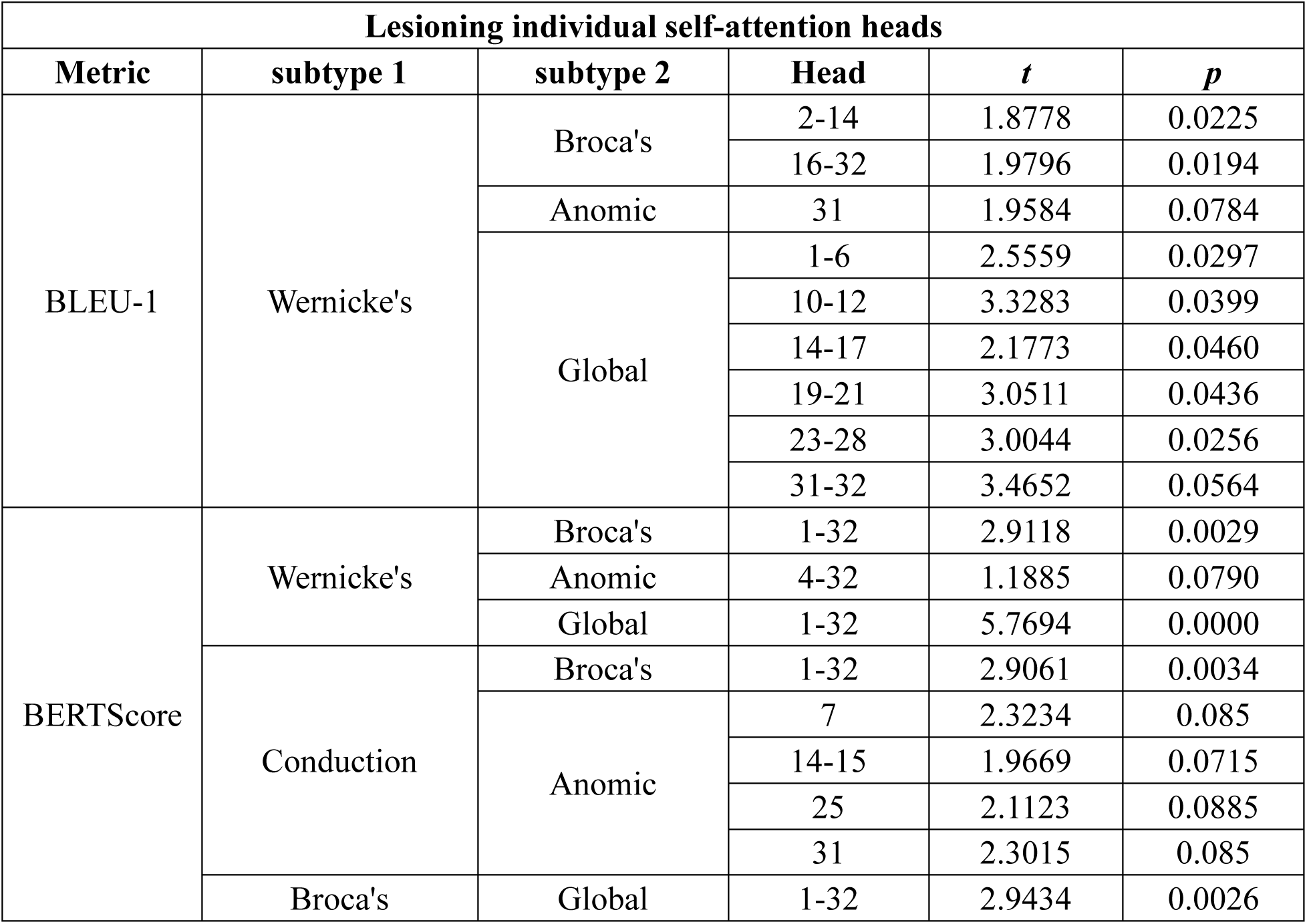
Results of non-parametric t-tests assessing the model’s similarity to various aphasia subtypes on the “Cookie Theft” task following lesions to individual self-attention heads.

**Supplementary table 6.**
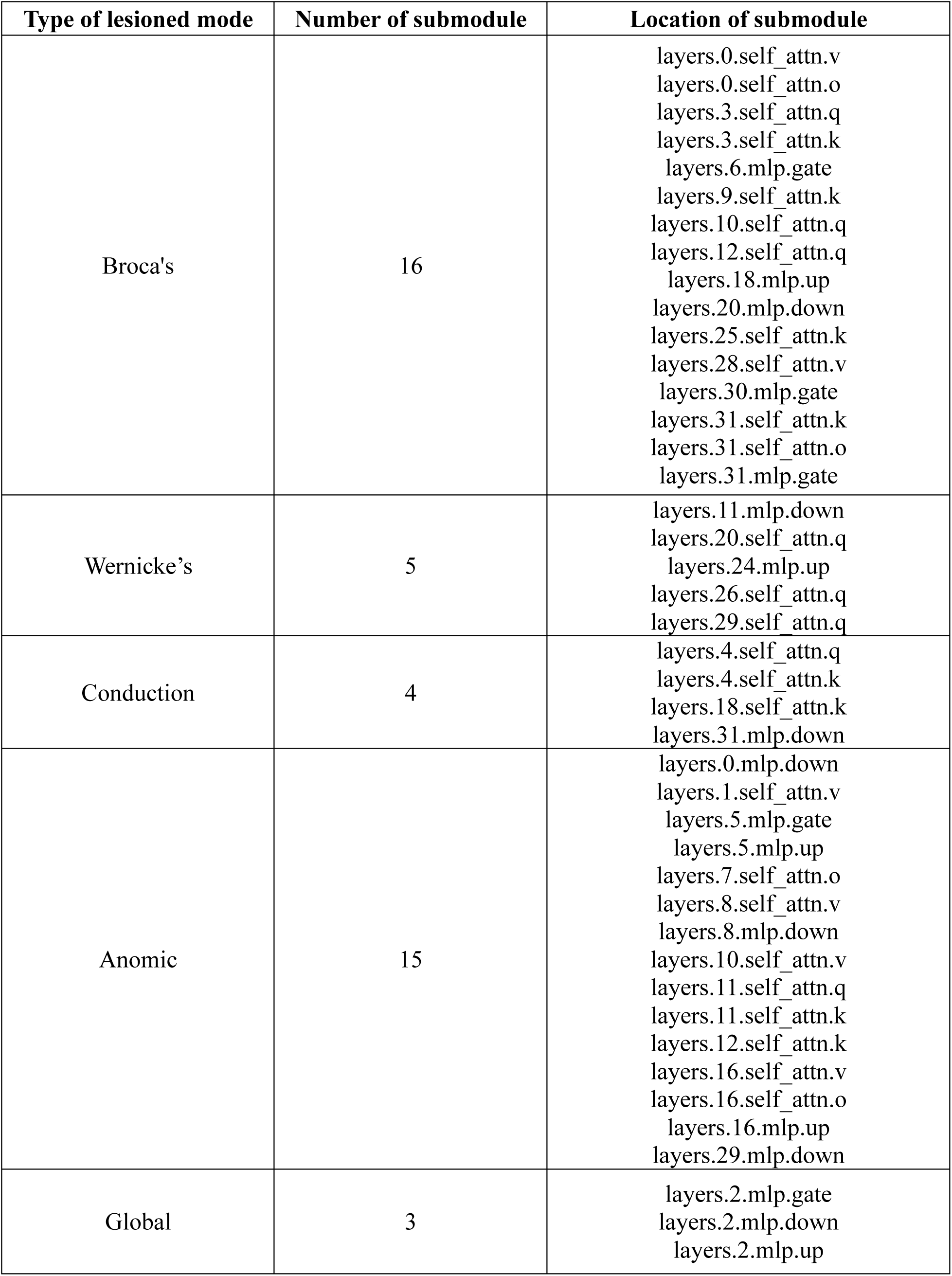
Distribution of parameters identified as critical for each aphasia behavior across submodules.

**Supplementary Table 7.** Model performance after lesioning each identified cluster on the “Cookie Theft” task.

| Model | Output 1 | Output 2 | Output 3 | Output 4 | Output 5 |
| --- | --- | --- | --- | --- | --- |
| Broca's | Details of the image include color, shape, details, color, shape, color, shape. | Details of the image include color. Shape. Size. Material. | Details of the image include sink. | Color used to describe the image, e.g., the details of this image. | Details of the image include water details, such as water shape. pH value. pH value. pH value. pH value. |
| Wernicke's | This picture depicts a family engaged in daily activities in the kitchen. The father is standing by the sink washing dishes; the mother is standing nearby, watching the children make paper decorations on the window. The children appear to be two boys, one sitting on a chair and the other standing to the right of the sink. Other items in the kitchen include cups, bottles, and bowls. | This picture shows a woman and two young boys standing in the kitchen, possibly washing dishes. The woman is wearing a blue shirt and black pants, while the two boys are dressed in white shirts and blue pants. The woman is holding a bowl, and the two boys are holding a knife and a spoon, respectively. There are several cups, bowls, plates, and bottles in the kitchen, some near the sink. | This image shows a mother and her children in the kitchen. She is standing by the sink, while her two children are on a ladder next to the sink. The girl is wearing pink clothes and holding a bowl, while the boy, who is not wearing a shirt, is holding a cup. Additionally, there are several bottles and bowls near the countertop. | This picture depicts a woman standing in the kitchen with two children beside her. The woman and children are dressed and holding some items. The woman's hair is tied into a ponytail, while the children are wearing hats. Several bottles and bowls are placed on the counter in front of the woman, and the children are standing to her left and right. | This picture shows a mother and two children playing in the kitchen. The mother is standing by the sink, with the children standing behind her. One boy is holding a small knife, while the other stands nearby. The mother has brown hair, bright eyes, and is smiling at the children. |
| Conduction | This picture shows a family scene, a woman and two kids in the kitchen. The woman is standing in the kitchen, and the kids are at the sink, the kid is at the sink, the kid is at the sink (repeatedly mentioned: “the kids are at the sink...”). | This picture shows a woman and two children in the kitchen. The woman is standing in the kitchen, and the children are beside her. The woman is wearing clothes, and the children are wearing clothes... (repeated “woman is wearing clothes, children are wearing clothes”). | This picture shows a woman and two children in the kitchen. The woman is standing in the kitchen, and the children are beside her. The woman is wearing clothes, and the children are wearing clothes... (repeated “woman is wearing clothes, children are wearing clothes”). | This picture shows a woman and two children in the kitchen. The woman is standing by the stove, and the children are sitting next to her. | This picture shows a woman and two children in the kitchen. The woman is standing in the kitchen, and the children are beside her... (repeated “woman is standing in the kitchen, children are beside her”). |
| Anomic | This picture depicts a woman and two children in the kitchen. They seem focused on their tasks, possibly washing dishes or cooking. The woman is standing by the sink, holding a spoon. One boy is sitting on a stool, and the other boy is standing near the sink. They seem engrossed in their work, oblivious to their surroundings. | This picture depicts a woman in the kitchen, standing by the sink and holding a bowl in her hand. She seems to be pouring water into the bowl, possibly preparing food. Behind her is a child standing by the wall near the sink, using his foot to draw on the wall. | This picture depicts a woman standing in the kitchen with two children, a boy and a girl. They seem focused on their tasks, possibly cooking or preparing food. The woman is wearing a white dress and a white hat, holding a spoon. Her hair is loose and not styled. The boy and girl are dressed similarly, with their hair also loose and unstyled. They seem focused on<br>their tasks,<br>possibly<br>cooking or<br>preparing<br>food. | This image depicts a woman standing in front of the kitchen counter, holding a bowl and a spoon. She appears to be pouring something into the water, possibly cooking for someone. In the background, a boy is sitting on a chair, while another person is watching from the side. They are all doing different things but focused on their things. | This picture depicts a woman standing in the kitchen, washing something over the sink. She is wearing a white dress and holding a cup and a bowl, possibly containing food or drink. Water is flowing from the tap onto her hands and face. She seems focused on her task but looks a little surprised when she sees the water drops. |
| Global | -----<br>-----<br>----- | t-000}', AND,<br>anding-t-t'-<br>-----<br>-----<br>-----<br>-----<br>----- | Tor_t',<br>M1000human<br>'or'.or<br>, %, ', %, '<br>/2%', or', [or_<br>M',<br>Shee_or', ', _<br>G '%oror-<br>_5%_ %_ \|V<br>iewor<br>[%_ 2%<br>%_ %_]- %<br>Care | '_t-<br>, %, '.or', or[ ', ', ', ',<br>, / ' She', '.or' ',<br>[' ', oror She', %', '.\|',<br>oning [human', _<br>ben She She% | 10_human', or[<br>[' ',<br>M00', %, 'Tor<br>or2%', ', ', ', %, '<br>% %', 'e'. _or<br>[' ', _<br>Careing.or<br>mor% [oror<br>[or_or_ |

**Supplementary Table 8.** Behavioral results of participants and models on the comprehension, production and repetition tasks.

| Accuracy scores of participants |  |  |  |
| --- | --- | --- | --- |
| Human | word associate matching | oral picture naming | oral word repetition |
| Control | 0.96±0.03 | 0.96±0.03 | 0.99±0.03 |
| Broca's | 0.92±0.05 | 0.75±0.15 | 0.85±0.18 |
| Wernicke's | 0.93±0.04 | 0.91±0.05 | 0.96±0.04 |
| Conduction | 0.95±0.04 | 0.93±0.06 | 0.82±0.13 |
| Anomic | 0.9±0.11 | 0.76±0.25 | 0.98±0.04 |
| Global | 0.83±0.1 | 0.44±0.22 | 0.62±0.23 |

| Accuracy scores of VisualCLA |  |  |  |
| --- | --- | --- | --- |
| Model | word associate matching | oral picture naming | oral word repetition |
| Intact | 0.83 | 0.9 | 1 |
| Broca's | 0.26 | 0.18 | 0.08 |
| Wernicke's | 0.78 | 0.84 | 0.83 |
| Conduction | 0.84 | 0.90 | 0.58 |
| Anomic | 0.60 | 0.64 | 0.75 |
| Global | 0.00 | 0.00 | 0.00 |

**Supplementary table 9.** Results of permutation *t*-test for inter-cluster connectivity.

| Model | Submodule 1 | Submodule 2 | Pearson' *r* | *p* |
| --- | --- | --- | --- | --- |
| Broca's | layers.0.self_attn.v | layers.0.self_attn.o | 0.9994 | 0.0000 |
|  | layers.0.self_attn.v | layers.10.self_attn.q | 0.1960 | 0.0000 |
|  | layers.0.self_attn.v | layers.12.self_attn.q | 0.1534 | 0.0065 |
|  | layers.0.self_attn.v | layers.31.self_attn.k | 0.2265 | 0.0035 |
|  | layers.0.self_attn.o | layers.10.self_attn.q | 0.2017 | 0.0000 |
|  | layers.0.self_attn.o | layers.12.self_attn.q | 0.1564 | 0.0063 |
|  | layers.0.self_attn.o | layers.31.self_attn.k | 0.2256 | 0.0049 |
|  | layers.3.self_attn.q | layers.3.self_attn.k | 0.9928 | 0.0000 |
|  | layers.3.self_attn.q | layers.6.mlp.gate | 0.5048 | 0.0000 |
|  | layers.3.self_attn.q | layers.9.self_attn.k | 0.5848 | 0.0000 |
|  | layers.3.self_attn.q | layers.10.self_attn.q | 0.6243 | 0.0000 |
|  | layers.3.self_attn.q | layers.12.self_attn.q | 0.5267 | 0.0000 |
|  | layers.3.self_attn.q | layers.25.self_attn.k | 0.2106 | 0.0000 |
|  | layers.3.self_attn.q | layers.31.self_attn.k | 0.3380 | 0.0000 |
|  | layers.3.self_attn.k | layers.6.mlp.gate | 0.4821 | 0.0000 |
|  | layers.3.self_attn.k | layers.9.self_attn.k | 0.6169 | 0.0000 |
|  | layers.3.self_attn.k | layers.10.self_attn.q | 0.6539 | 0.0000 |
|  | layers.3.self_attn.k | layers.12.self_attn.q | 0.5621 | 0.0000 |
|  | layers.3.self_attn.k | layers.25.self_attn.k | 0.2230 | 0.0000 |
|  | layers.3.self_attn.k | layers.31.self_attn.k | 0.3617 | 0.0000 |
|  | layers.6.mlp.gate | layers.9.self_attn.k | 0.1517 | 0.0054 |
|  | layers.6.mlp.gate | layers.18.mlp.up | 0.6918 | 0.0000 |
|  | layers.6.mlp.gate | layers.20.mlp.down | 0.2246 | 0.0000 |
|  | layers.6.mlp.gate | layers.25.self_attn.k | 0.2017 | 0.0000 |
|  | layers.6.mlp.gate | layers.28.self_attn.v | 0.2383 | 0.0000 |
|  | layers.6.mlp.gate | layers.30.mlp.gate | 0.2273 | 0.0001 |
|  | layers.6.mlp.gate | layers.31.self_attn.o | 0.2371 | 0.0000 |
|  | layers.6.mlp.gate | layers.31.mlp.gate | 0.2342 | 0.0000 |
|  | layers.9.self_attn.k | layers.10.self_attn.q | 0.8826 | 0.0000 |
|  | layers.9.self_attn.k | layers.12.self_attn.q | 0.8934 | 0.0000 |
|  | layers.9.self_attn.k | layers.25.self_attn.k | 0.5172 | 0.0000 |
|  | layers.9.self_attn.k | layers.31.self_attn.k | 0.6404 | 0.0000 |
|  | layers.10.self_attn.q | layers.12.self_attn.q | 0.9199 | 0.0000 |
|  | layers.10.self_attn.q | layers.25.self_attn.k | 0.4821 | 0.0000 |
|  | layers.10.self_attn.q | layers.31.self_attn.k | 0.6593 | 0.0000 |
|  | layers.12.self_attn.q | layers.25.self_attn.k | 0.5300 | 0.0000 |
|  | layers.12.self_attn.q | layers.31.self_attn.k | 0.6960 | 0.0000 |
|  | layers.18.mlp.up | layers.20.mlp.down | 0.6711 | 0.0000 |
|  | layers.18.mlp.up | layers.28.self_attn.v | 0.6764 | 0.0000 |
|  | layers.18.mlp.up | layers.30.mlp.gate | 0.6672 | 0.0000 |
|  | layers.18.mlp.up | layers.31.self_attn.o | 0.6733 | 0.0000 |
|  | layers.18.mlp.up | layers.31.mlp.gate | 0.6715 | 0.0000 |
|  | layers.20.mlp.down | layers.28.self_attn.v | 0.9995 | 0.0000 |
|  | layers.20.mlp.down | layers.30.mlp.gate | 0.9996 | 0.0000 |
|  | layers.20.mlp.down | layers.31.self_attn.o | 0.9994 | 0.0000 |
|  | layers.20.mlp.down | layers.31.mlp.gate | 0.9995 | 0.0000 |
|  | layers.25.self_attn.k | layers.31.self_attn.k | 0.4968 | 0.0000 |
|  | layers.28.self_attn.v | layers.30.mlp.gate | 0.9997 | 0.0000 |
|  | layers.28.self_attn.v | layers.31.self_attn.o | 0.9998 | 0.0000 |
|  | layers.28.self_attn.v | layers.31.mlp.gate | 0.9998 | 0.0000 |
|  | layers.30.mlp.gate | layers.31.self_attn.o | 0.9997 | 0.0000 |
|  | layers.30.mlp.gate | layers.31.mlp.gate | 0.9998 | 0.0000 |
|  | layers.31.self_attn.o | layers.31.mlp.gate | 1.0000 | 0.0000 |
| Wernicke's | layers.11.mlp.down | layers.24.mlp.up | 0.6412 | 0.0000 |
|  | layers.20.self_attn.q | layers.26.self_attn.q | 0.8111 | 0.0000 |
|  | layers.20.self_attn.q | layers.29.self_attn.q | 0.6109 | 0.0000 |
|  | layers.26.self_attn.q | layers.29.self_attn.q | 0.5315 | 0.0000 |
| Conduction | layers.4.self_attn.q | layers.4.self_attn.k | 0.9943 | 0.0000 |
|  | layers.4.self_attn.q | layers.18.self_attn.k | 0.6126 | 0.0000 |
|  | layers.4.self_attn.k | layers.18.self_attn.k | 0.6421 | 0.0000 |
| Anomic | layers.0.mlp.down | layers.11.self_attn.q | 0.2523 | 0.0000 |
|  | layers.0.mlp.down | layers.11.self_attn.k | 0.2670 | 0.0000 |
|  | layers.0.mlp.down | layers.12.self_attn.k | 0.3002 | 0.0000 |
|  | layers.1.self_attn.v | layers.5.mlp.up | 0.1112 | 0.0400 |
|  | layers.1.self_attn.v | layers.7.self_attn.o | 0.9321 | 0.0000 |
|  | layers.1.self_attn.v | layers.8.self_attn.v | 0.9152 | 0.0000 |
|  | layers.1.self_attn.v | layers.8.mlp.down | 0.6292 | 0.0000 |
|  | layers.1.self_attn.v | layers.10.self_attn.v | 0.9318 | 0.0000 |
|  | layers.1.self_attn.v | layers.16.self_attn.v | 0.9398 | 0.0000 |
|  | layers.1.self_attn.v | layers.16.self_attn.o | 0.9435 | 0.0000 |
|  | layers.1.self_attn.v | layers.16.mlp.up | 0.6347 | 0.0000 |
|  | layers.1.self_attn.v | layers.29.mlp.down | 0.9413 | 0.0000 |
|  | layers.5.mlp.gate | layers.5.mlp.up | 0.9921 | 0.0000 |
|  | layers.5.mlp.gate | layers.7.self_attn.o | 0.1115 | 0.0372 |
|  | layers.5.mlp.gate | layers.8.self_attn.v | 0.1818 | 0.0008 |
|  | layers.5.mlp.gate | layers.8.mlp.down | 0.7185 | 0.0000 |
|  | layers.5.mlp.gate | layers.11.self_attn.q | 0.2809 | 0.0000 |
|  | layers.5.mlp.gate | layers.11.self_attn.k | 0.2579 | 0.0000 |
|  | layers.5.mlp.gate | layers.12.self_attn.k | 0.1506 | 0.0046 |
|  | layers.5.mlp.gate | layers.16.mlp.up | 0.6288 | 0.0000 |
|  | layers.5.mlp.up | layers.7.self_attn.o | 0.1552 | 0.0059 |
|  | layers.5.mlp.up | layers.8.self_attn.v | 0.2244 | 0.0001 |
|  | layers.5.mlp.up | layers.8.mlp.down | 0.7419 | 0.0000 |
|  | layers.5.mlp.up | layers.11.self_attn.q | 0.2388 | 0.0000 |
|  | layers.5.mlp.up | layers.11.self_attn.k | 0.2157 | 0.0000 |
|  | layers.5.mlp.up | layers.12.self_attn.k | 0.1124 | 0.0426 |
|  | layers.5.mlp.up | layers.16.mlp.up | 0.6535 | 0.0000 |
|  | layers.7.self_attn.o | layers.8.self_attn.v | 0.9923 | 0.0000 |
|  | layers.7.self_attn.o | layers.8.mlp.down | 0.7072 | 0.0000 |
|  | layers.7.self_attn.o | layers.10.self_attn.v | 0.9976 | 0.0000 |
|  | layers.7.self_attn.o | layers.16.self_attn.v | 0.9958 | 0.0000 |
|  | layers.7.self_attn.o | layers.16.self_attn.o | 0.9956 | 0.0000 |
|  | layers.7.self_attn.o | layers.16.mlp.up | 0.6827 | 0.0000 |
|  | layers.7.self_attn.o | layers.29.mlp.down | 0.9955 | 0.0000 |
|  | layers.8.self_attn.v | layers.8.mlp.down | 0.7462 | 0.0000 |
|  | layers.8.self_attn.v | layers.10.self_attn.v | 0.9880 | 0.0000 |
|  | layers.8.self_attn.v | layers.16.self_attn.v | 0.9817 | 0.0000 |
|  | layers.8.self_attn.v | layers.16.self_attn.o | 0.9810 | 0.0000 |
|  | layers.8.self_attn.v | layers.16.mlp.up | 0.7083 | 0.0000 |
|  | layers.8.self_attn.v | layers.29.mlp.down | 0.9809 | 0.0000 |
|  | layers.8.mlp.down | layers.10.self_attn.v | 0.6742 | 0.0000 |
|  | layers.8.mlp.down | layers.16.self_attn.v | 0.6602 | 0.0000 |
|  | layers.8.mlp.down | layers.16.self_attn.o | 0.6646 | 0.0000 |
|  | layers.8.mlp.down | layers.16.mlp.up | 0.9429 | 0.0000 |
|  | layers.8.mlp.down | layers.29.mlp.down | 0.6589 | 0.0000 |
|  | layers.10.self_attn.v | layers.16.self_attn.v | 0.9981 | 0.0000 |
|  | layers.10.self_attn.v | layers.16.self_attn.o | 0.9975 | 0.0000 |
|  | layers.10.self_attn.v | layers.16.mlp.up | 0.6557 | 0.0000 |
|  | layers.10.self_attn.v | layers.29.mlp.down | 0.9978 | 0.0000 |
|  | layers.11.self_attn.q | layers.11.self_attn.k | 0.9953 | 0.0000 |
|  | layers.11.self_attn.q | layers.12.self_attn.k | 0.9230 | 0.0000 |
|  | layers.11.self_attn.k | layers.12.self_attn.k | 0.9247 | 0.0000 |
|  | layers.16.self_attn.v | layers.16.self_attn.o | 0.9997 | 0.0000 |
|  | layers.16.self_attn.v | layers.16.mlp.up | 0.6504 | 0.0000 |
|  | layers.16.self_attn.v | layers.29.mlp.down | 0.9997 | 0.0000 |
|  | layers.16.self_attn.o | layers.16.mlp.up | 0.6544 | 0.0000 |
|  | layers.16.self_attn.o | layers.29.mlp.down | 0.9999 | 0.0000 |
|  | layers.16.mlp.up | layers.29.mlp.down | 0.6475 | 0.0000 |
| Global | layers.2.mlp.gate | layers.2.mlp.down | 0.9962 | 0.0000 |
|  | layers.2.mlp.gate | layers.2.mlp.up | 0.9994 | 0.0000 |
|  | layers.2.mlp.down | layers.2.mlp.up | 0.9975 | 0.0000 |

